# Early events marking lung fibroblast transition to profibrotic state in idiopathic pulmonary fibrosis

**DOI:** 10.1101/2022.07.29.501956

**Authors:** Minxue Jia, Lorena Rosas, Maria G. Kapetanaki, Tracy Tabib, John Sebrat, Tamara Cruz, Anna Bondonese, Ana L. Mora, Robert Lafyatis, Mauricio Rojas, Panayiotis .V. Benos

## Abstract

**Background:** Idiopathic Pulmonary Fibrosis (IPF) is an age-associated progressive lung disease with accumulation of scar tissue impairing gas exchange. Previous high-throughput studies elucidated the role of cellular heterogeneity and molecular pathways in advanced disease. However, critical pathogenic pathways occurring in the transition of fibroblasts from normal to profibrotic have been largely overlooked.

**Methods:** We used single cell transcriptomics (scRNA-seq) from lungs of healthy controls and IPF patients (lower and upper lobes). We identified fibroblast subclusters, genes and pathways associated to early disease. Immunofluorescence assays validated the role of MOXD1 early in fibrosis.

**Findings:** We identified four distinct fibroblast subgroups, including one marking the normal-to-profibrotic state transition. Our results show for the first time that global downregulation of ribosomal proteins and significant upregulation of the majority of copper-binding proteins, including MOXD1, mark the IPF transition. We find no significant differences in gene expression in IPF upper and lower lobe samples, which were selected to have low and high degree of fibrosis, respectively.

**Interpretation:** Early events during IPF onset in fibroblasts include dysregulation of ribosomal and copper-binding proteins. Fibroblasts in early stage IPF have already acquired a profibrotic phenotype while hallmarks of advanced disease, including fibroblast foci and honeycomb formation, are still not evident. The new transitional fibroblasts we discover could prove very important for studying the role of fibroblast plasticity in disease progression and help develop early diagnosis tools and therapeutic interventions targeting earlier disease states.

## 1. Introduction

Idiopathic Pulmonary Fibrosis (IPF) is an age-dependent chronic lung disease affecting individuals generally over 60 years old (1). The mechanisms driving the disease development and progression are still not fully understood (2). Gene-by-gene analysis and high-throughput studies have promoted the field over the years, offering valuable insights into the pathophysiology of the disease (3-5). Similar to other diseases, single cell approaches have the potential to advance this knowledge even further by fully dissecting the disease milieu and allowing investigators to assess individualized cell contributions and complex cellular dynamics during disease emergence and progression (6-8).

Extensive research has provided valuable insight into the origin and contribution of fibroblasts and especially myofibroblasts in pulmonary fibrosis (9, 10). All evidence supports the hypothesis that these are heterogeneous groups of cells, undergoing very distinct transition processes dictated by the disease microenvironments (9-11). They share common characteristics and express a set of biomolecules that drive fibrosis (12). Single cell transcriptomics allow dissection of this heterogeneous population. Murine models have been used to better understand how fibroblast subtypes contribute to fibrosis (9). More recently, single cell RNA sequencing (scRNA-seq) of fresh human tissue revealed cell-specific differences between normal and fibrotic tissue (8, 13).

To study the disease onset and progression, our group has performed scRNA-seq on upper and lower lobes of fresh human explanted IPF lungs and on healthy controls, identifying a subpopulation of proliferating SPP1^Hi^ macrophages with a potential role in lung fibrosis (8). Here, we reanalyze the raw sequencing data after performing an imputation step and focus on the early events that could drive transition of normal lung fibroblasts to profibrotic. Our analyses reveal four major fibroblast clusters showing unique characteristics regarding their gene expression and related to their tissue of origin (control or IPF). A closer look at each cluster confirms previously reported changes associated with IPF and -more importantly-reveals a profibrotic state of the upper (unaffected) lung. Gene expression patterns in each cluster reveal an expected dysregulation of fibrosis-associated genes. A novel finding is the dysregulation of genes coding for copper-binding proteins during both early and late stages of the disease. Immunohistochemistry assays in IPF lung fibroblasts show high levels of one of the top differentially expressed copper-binding genes, MOXD1. Furthermore, pseudotime analysis identifies a distinct group of fibroblasts in the process of acquiring a profibrotic phenotype while undergoing a global downregulation of genes coding for ribosomal proteins. Both copper-binding (14) and ribosomal protein pathways (15-17) have been previously associated to senescence. The discovery of this new type of transitional fibroblasts is important as it provides insights into the early events of fibrosis and may prove more helpful in finding suitable targets for early diagnosis and possible therapeutic interventions.

## 2. Methods

### 2.1. Single cell RNA sequencing (scRNA-seq)

Raw scRNA-seq data were derived from our previous publication (8). Briefly, normal and IPF lung tissue (**Table S1**) was obtained and processed to obtain single cell suspension. Single cell libraries were prepared using the 10X Genomics Chromium instrument and V2 chemistry. Sequencing was performed on an Illumina NextSeq-500 instrument.

### 2.2. scRNA-seq Data Analysis

scRNA-seq raw count and cell-UMI (Unique Molecular Identifier) count matrix were generated by Cell Ranger (8). Single-cell Analysis Via Expression Recovery (Saver) was used to impute dropout events in gene expression (18). Seurat (version 2.3.4) was used to normalize gene expression, perform differentially expressed gene analysis, identify distinct cell populations and visualize clusters graphically (19, 20). The cell-UMI matrix was filtered and only cells expressing at least 200 genes were further analyzed. Cells containing greater than 35% of mitochondrial genes were also excluded from the analysis. Highly variable genes were identified, based on their average expression and dispersion, and would be used in the downstream analysis. Data were scaled and the number of UMIs per cell as well as the percentage of mitochondrial gene content were regressed out. In this study, we further removed the effect from technical or biological confounders using *Harmony*, which integrates multiple scRNA seq datasets by projecting cells into a shared embedding in which cells are grouped by cell type, not the specific conditions related to the datasets (21). t-distributed stochastic neighbor embedding (t-SNE) plots based on Harmony embeddings were generated to assign clusters and each cluster was identified by differentially expressed genomic signatures.

### 2.3. Analysis of Fibroblast cluster from scRNA-seq data

The fibroblast cells from scRNA-seq data were processed by Seurat and Harmony to identify subclusters of fibroblast cells. Destiny was used for pseudotime analysis of the subclusters (20). Velocyto was used to estimate the time derivative of the gene expression state (22). RNA velocity was estimated using gene-relative model with k-nearest neighbor cell pooling (k=20) and velocity fields were projected into a UMAP-based embedding through SeuratWrappers and Seurat (version 3.1.0). We also used AddModuleScore of Seurat to calculate average expression level of i) copper binding (23), ii) senescence (24) and iii) ribosome biogenesis (Gene Ontology Browser) gene sets on a single cell level, normalized by randomly selected control feature set.

### 2.4. Immunofluorescence

Fresh lung tissue (**Table S1**) was fixed with 4% paraformaldehyde and embedded in OCT (Tissue-Tek® Sakura® Finetek, US). Cultured fibroblasts (**Table S1**) were plated in chamber-slides and fixed with 2% paraformaldehyde. OCT sections (5μM thick) and fixed fibroblasts were stained with the primary/secondary antibodies listed in the **Table S2** (Supplemental data). Slides were mounted using ProLong Gold Antifade Mountant with DAPI (Life technologies, USA). Images were obtained using an Olympus Fluoview 1000-3 Confocal Microscope (20x objective).

## 3. Results

### 3.1. Cell types in IPF and healthy lungs

A mixed population of cells (48,023 cells) from the lungs of three healthy control (21,485 cells) and six IPF (26,538 cells) samples (from tree donors) was sequenced using next generation single cell sequencing. An important study design includes tissue from three upper lobes (IPF_UP; 12,577 cells) and fibrotic tissue from three lower lobes (IPF_LOW; 13,961 cells) (**Figure 1A**). In general, samples from the upper lobes presented a relatively unaffected lung histology, while all samples from the lower lobes showed advanced fibrosis (8). Here, we performed imputation of dropouts and gene expression profiles were clustered and visualized (t-SNE). Our analysis resulted in 20 clusters (**Figure 1B**) that were identified as distinct cell types (**Figure 1C**), using previously described cell markers (8). Each cluster had variable cell counts coming from healthy and IPF upper and lower lung lobes (**Figure 1D-E** and **S1)**

**Figure 1.**
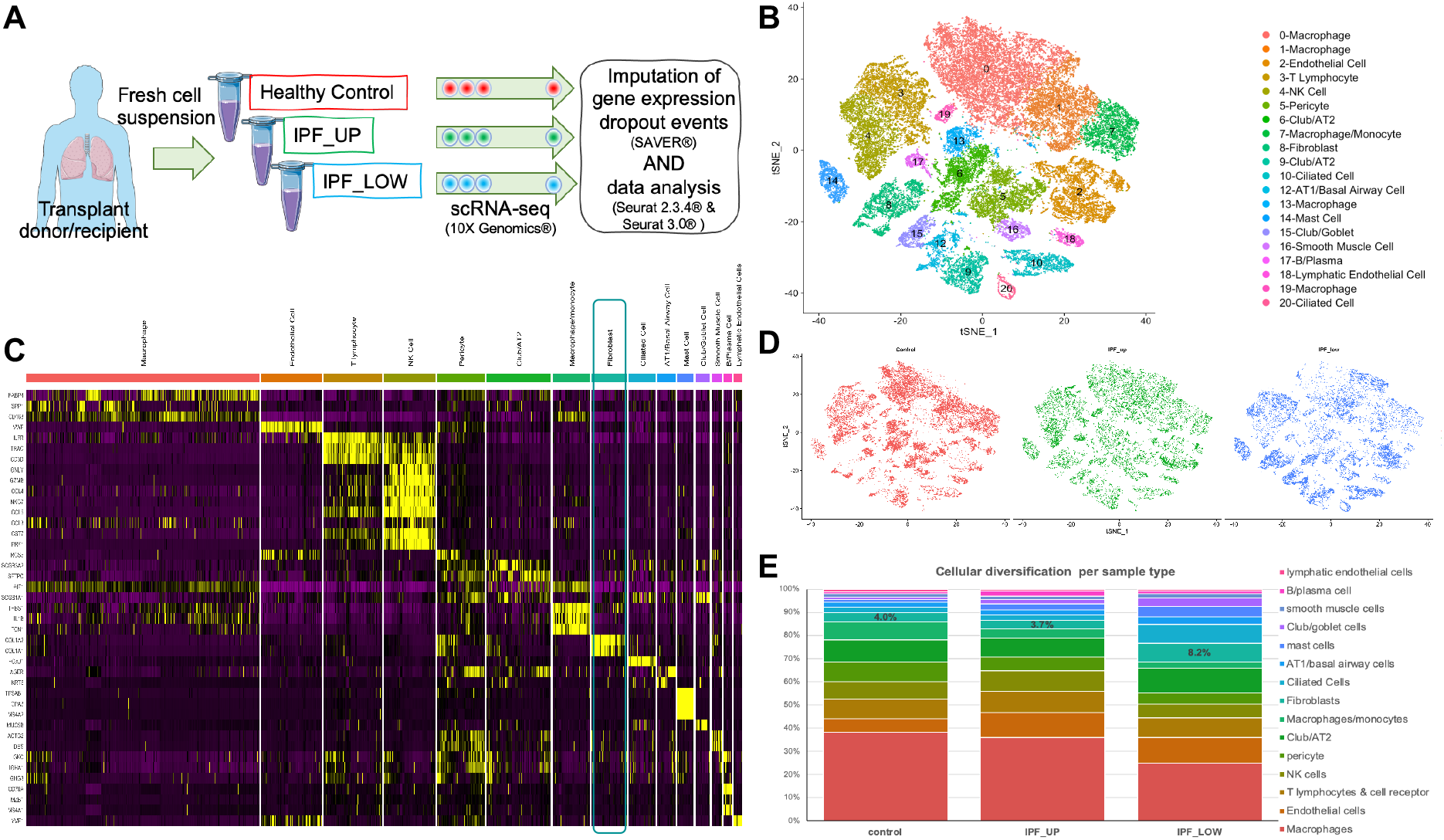
Analysis of scRNA-seq data from control and IPF lung cells identifies a complex network of cell types. **(A)** A suspension of single cells was generated from lung explants of control or IPF patients and was subsequently sequenced using the 10X genomics scRNA platform. **(B)** t-SNE plot of cell clustering after data imputation (raw sequencing data: Morse *et al*.) **(C)** Expression heatmap of genes that are used to identify cell type in each cluster. **(D)** Cells in the t-SNE plots are colored according to their origin (control, IPF_UP or IPF_LOW). **(E)** Cellular composition of each sample type.

### 3.2. Fibroblast (FB) clusters’ characteristics

Fibroblasts (**Figure 1C**, green box) were re-clustered and visualized in two dimensions using t-SNE (**Figure 2A**). Overall, the clustering of fibroblasts from 449 control, 404 IPF_UP and 1085 IPF_LOW samples revealed four different groups. A fifth cluster, neighboring B-cells, was found to contain cells expressing HLA type II family genes and was excluded from this analysis (**Figure S2**). Despite our best efforts, one of the IPF upper lung lobe samples (SC154) showed extensive fibrosis compared to normal controls, posing a risk of masking any differences between the IPF_UP and IPF_LOW fibroblasts. We decided to include this sample in our analysis since it clustered similarly to the other IPF_UP samples (**Figure S3**).

**Figure 2.**
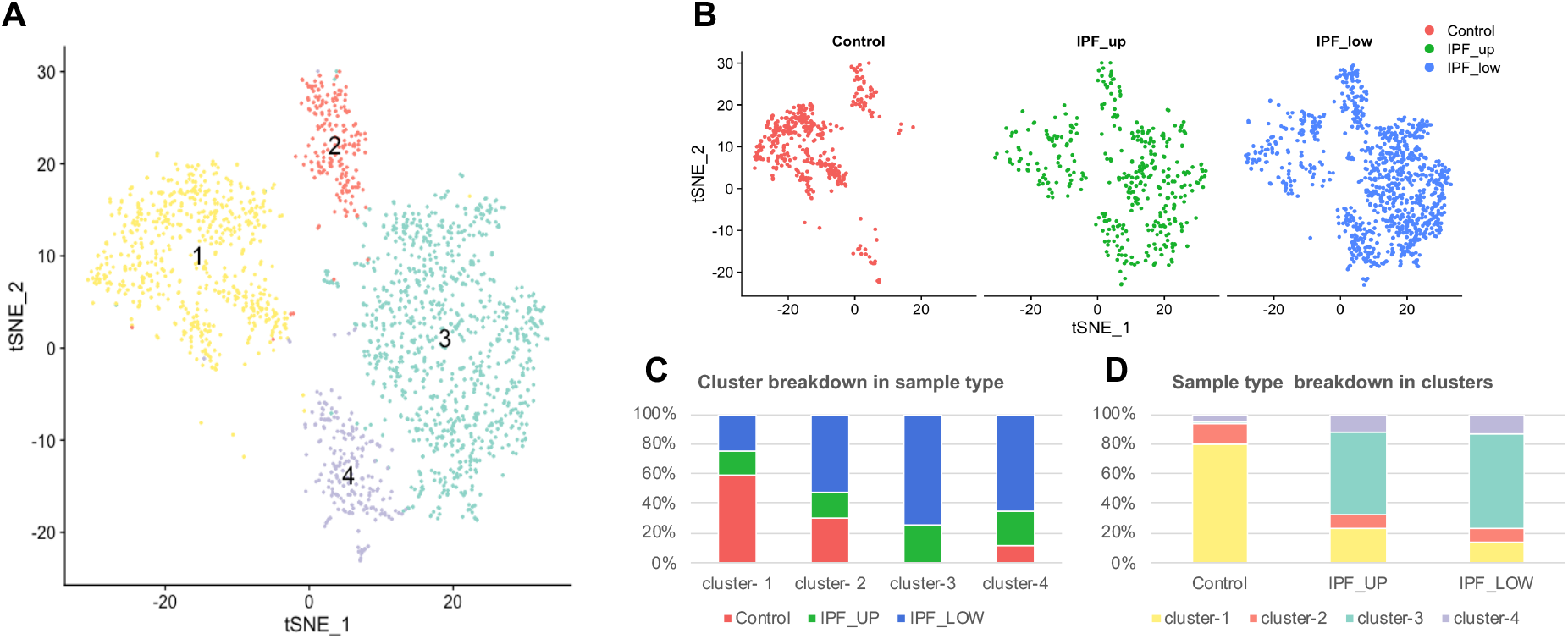
Analysis of scRNA sequencing data from control and IPF lung fibroblasts identifies four distinct clusters/subtypes. (**A**) t-SNE plot of fibroblast clusters (**B**) Cells in the tSNE plots are colored according to their origin (control, IPF_UP or IPF_LOW). (**C**) Contribution of each sample type to each cluster. (**D**) Partition of each cell origin into the four fibroblast clusters.

Cluster-1 (608 cells, 31% of all FB) contained mainly cells from the healthy control donors (80% of the control FBs were in Cluster-1) as well as a smaller percentage of the IPF_UP and IPF_LOW cells (24% and 14% respectively). Cluster-2, a much smaller cluster (197 cells, 10% of all FB), included cells from each sample origin. Cluster-3 (920 cells, 48% of all FB) contained mainly IPF_UP and IPF_LOW (56% and 64% respectively). Only 1% of the healthy control FBs classified as Cluster-3. Cluster-4 (213 cells, 11% of all FB) contained a slightly higher percentage of the IPF_UP and IPF_LOW (12% and 13% respectively) than the healthy control fibroblasts (6%). The breakdown of each cluster to sample types is summarized in **Figures 2B-C** while the breakdown of each sample to different clusters is summarized in **Figure 2D**.

Since fibroblasts in Cluster-1 came predominantly from control samples we consider this to represent the “normal state”; while those in Cluster-3, which came almost exclusively form IPF_UP and IPF_LOW, represent the “disease state”. Cluster-1, Cluster-2 and Cluster-4 had distinct expression profiles, but they were all present in healthy and IPF lungs (**Figure 2C**). Overall, IPF_UP and IPF_LOW fibroblasts showed similar clustering and compositional patterns (**Figure 2B** and **2D**). These results showed that fibroblasts from the relatively non-fibrotic upper lobes had similar molecular signatures to those from the highly fibrotic lower lobes, suggesting that molecular changes in lung fibroblasts precede the morphological changes identified by histological examination.

Differentially expressed genes in each cluster are listed in **Table S3** and expression of the top 16 genes in these lists is further described per cluster and sample type (**Figures 3-6**). Briefly, fibroblasts in Cluster-1 had higher expression in 263 genes (FDR<0.05) with the highest being: ICAM1, CXCL1, CXCL2, CXCL3, CXCL8, CCL2, IL6, PTX3, IER3, GADD45B, THBS1, SOD2 and NFKBIA which are associated with immune response and regulation of inflammation (**Figure 3** and **Table S4**).

**Figure 3.**
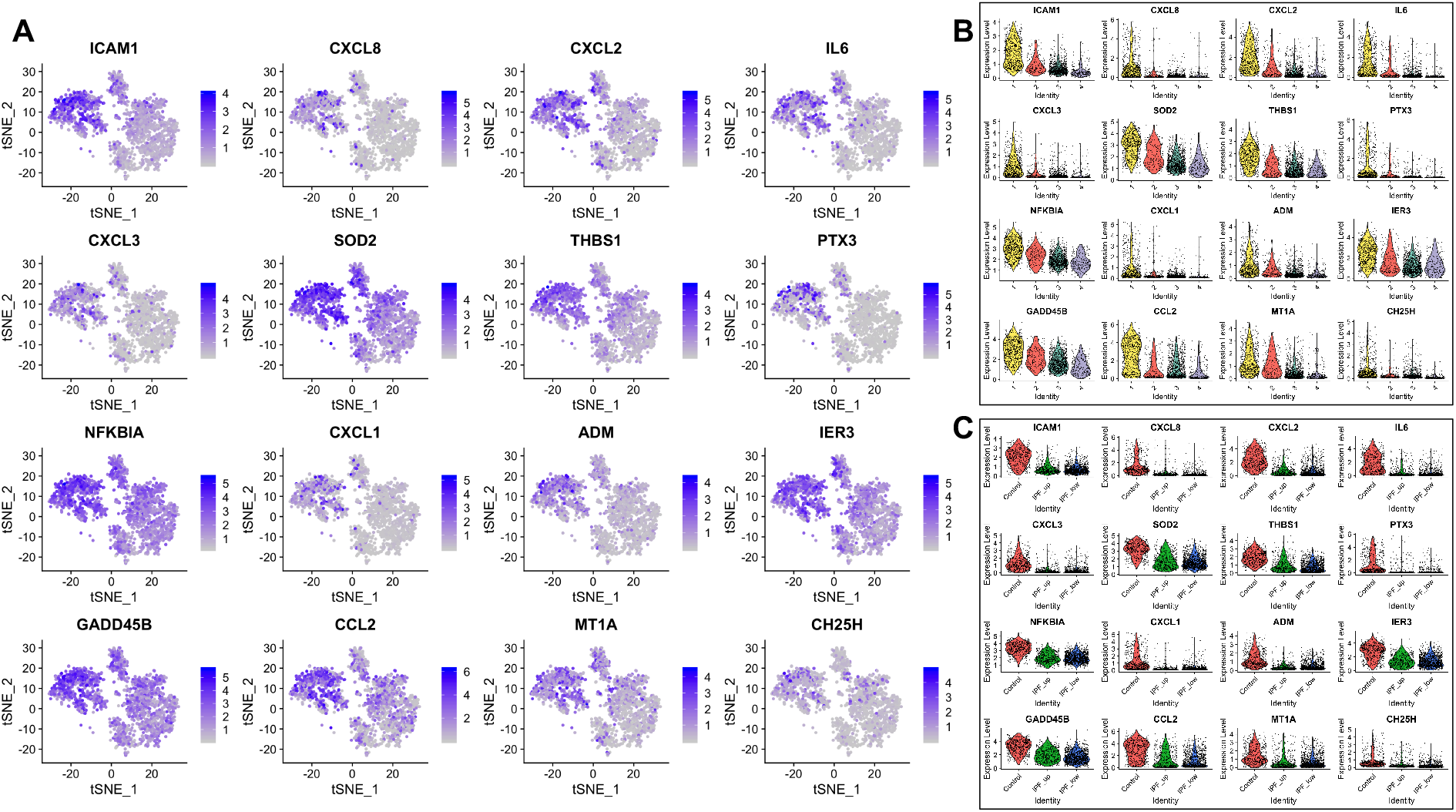
Top 16 most upregulated genes in Cluster-1. **(A)** tSNE plots. **(B)** Expression by cluster. **(C)** Expression by tissue of origin.

Among the 118 genes that were significantly upregulated in Cluster-2, CXCL14, SFRP2 and SFRP4 had higher expression levels in IPF samples. Interestingly, the highest levels of CXCL14 were detected in IPF_LOW fibroblasts, being one of the most significant differences observed overall between IPF_UP and IPF_LOW samples. Profibrotic gene CXCL14 is shown to play a regulatory role in immune response and inflammation while SFRP2 and SFRP4 are regulators (inhibitors) of Wnt signaling (**Figure 4** and **Table S4**).

**Figure 4.**
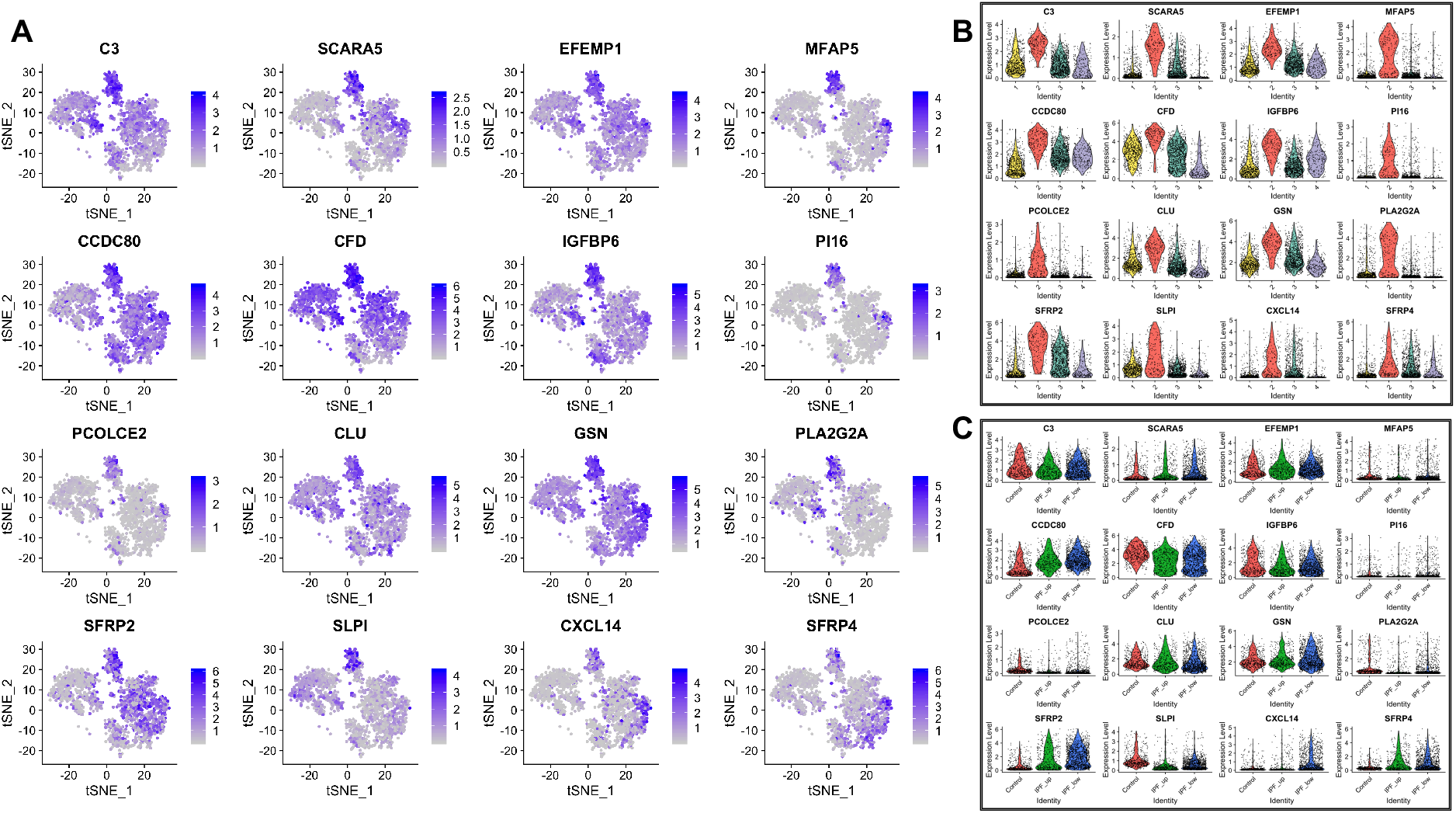
Top 16 most upregulated genes in Cluster-2. **(A) (B) (C)** as above.

The majority of the top 16 upregulated genes in Cluster-3 (out of the total 219 differentially expressed) have been previously shown to increase in lung fibrosis (**Figure 5** and **Table S4**). Our results corfirmed that cells expressing myofibroblasts markers like POSTN and ASPN also expressed LTBP2, LTBP1, BGN, DPT and the highest levels of COL3A1 and COL8A1. Interestingly, 68-93% of the cells on this cluster that had low expression of myofibroblast-associated genes also had high expression levels of IPF-associated profibrotic markers such as MFAP4, LTBP1, BGN, COMP, MMP2, COL3A1 and COL8A1. All 16 genes had similar expression in IPF_UP and IPF_LOW samples. CXCL14, which was upregulated in Cluster-2, was also upregulated in the Cluster-3 IPF_LOW compared to IPF_UP.

**Figure 5.**
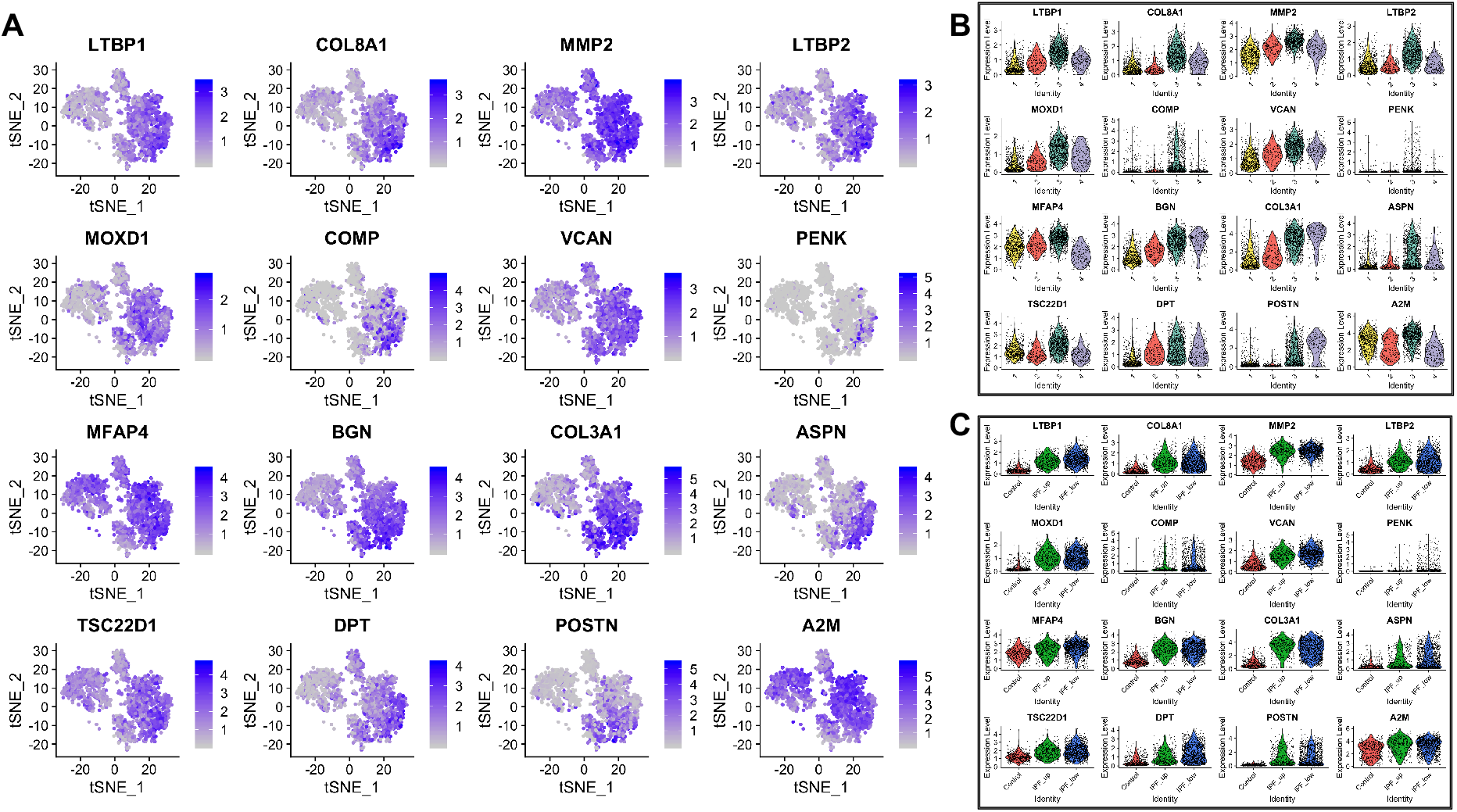
Top 16 most upregulated genes in Cluster-3. **(A) (B) (C)** as above.

Cells in Cluster-4 expressed profibrotic and myofibroblast markers like TNC, collagens, SPARC, POSTN, FN1 and TPM2 in higher levels than cells in Cluster-3. All 16 top upregulated genes in this group had similar expression levels in IPF_UP and IPF_LOW samples (**Figure 6** and **Table S4**).

**Figure 6.**
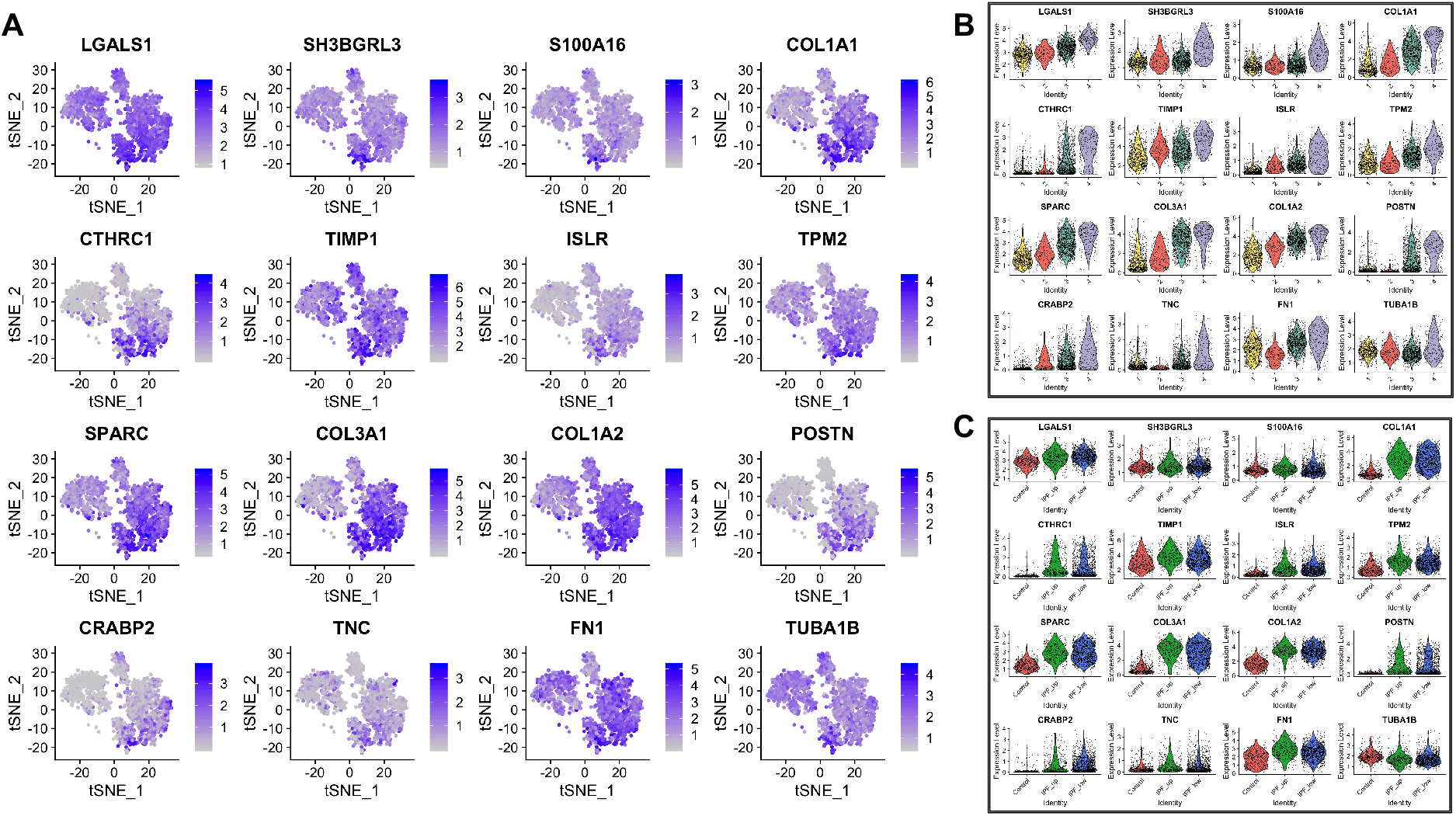
Top 16 most upregulated genes in Cluster-4. **(A) (B) (C)** as above.

Full names and relevant references for all genes mentioned above are listed in Supplemental Data **Table S4**.

### 3.3. Trajectory and RNA velocity Analysis reveals early disease-associated events

Single cell trajectory inference and pseudotime analysis are often used to study cellular dynamics or transitional states and special organization of cells in tissues (25). Each cell is assigned a numeric value (pseudotime) which indicates where in the underlying dynamic biological process that cell falls into. These methods allow the visualization of intermediate stages that connect distinct cell states, which are often overlooked during cell clustering. We used diffusion map (destiny) to perform pseudotime analysis on the four fibroblast subclusters (**Figure 7A**). The diffusion components, as measures of pseudotime (26), revealed temporal ordering and cellular decision on the single-cell transcriptome level. The diffusion map embedding of all fibroblasts resulted in a structure showing a progressive cell transition from Cluster-1 to 4 (**Figure 7A**, arrow). Interestingly, cells in Cluster-1 had two distinct subpopulations (**Figure 7A**, ovals) which corresponded to cells from Control and IPF samples with the latter being closer to Cluster-2 and 3 (**Figure 7B**). This suggests that IPF fibroblasts in Cluster-1, despite their overall “healthy” phenotype, have been influenced by the disease environment.

**Figure 7.**
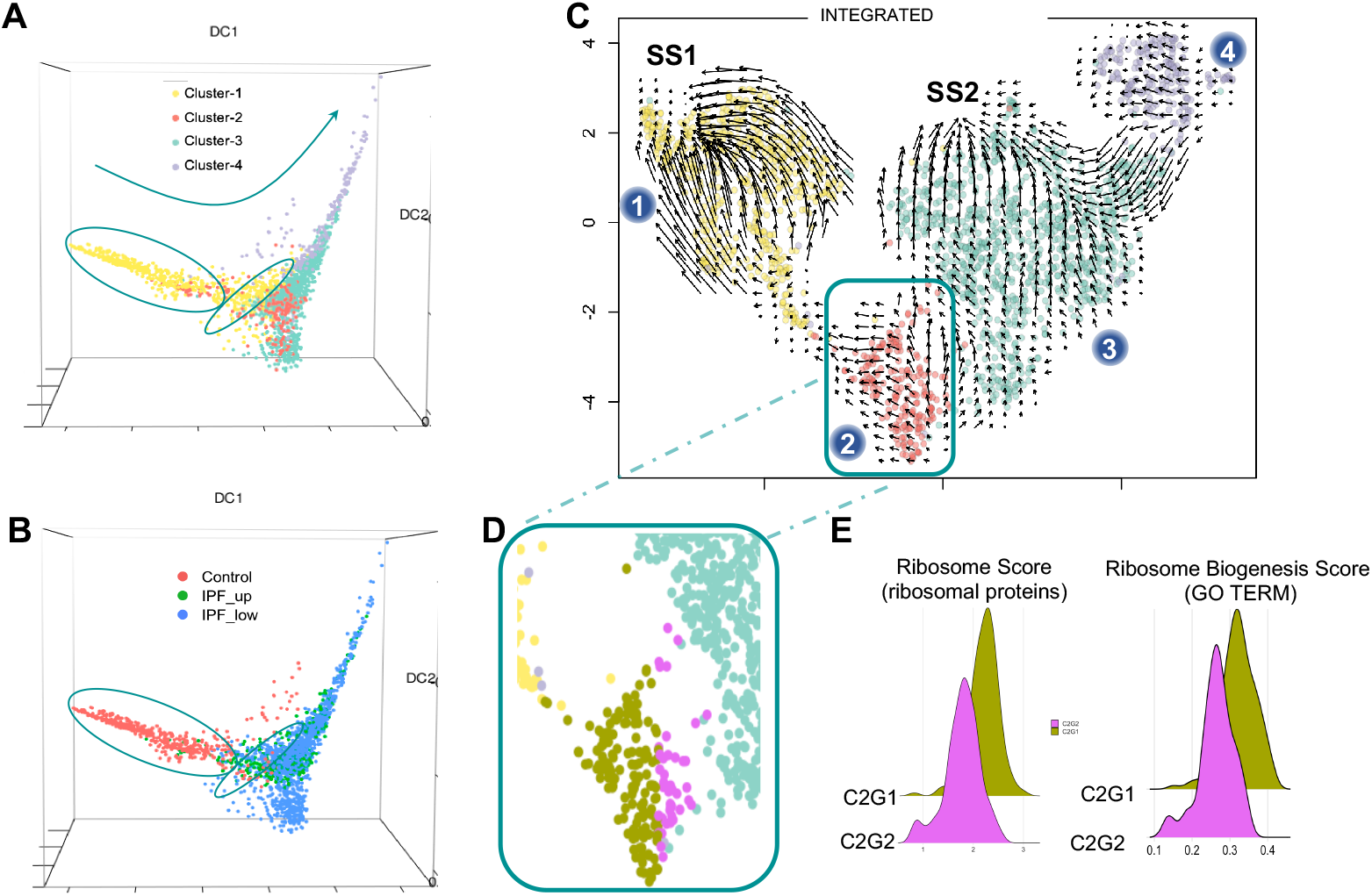
Pseudotime analysis of fibroblasts. RNA trajectory analysis shows (**A**) two distinct cell subgroups (ovals) in Cluster-1 (**B**) depending on the cell origin (control or IPF). Overall, the majority of the cells follow a pseudotime trajectory suggesting a phenotypic transition from Cluster-1 to Cluster-2 to Cluster-3 and finally Cluster-4 (arrow). (**C**) Velocity analysis of all fibroblasts suggests that Cluster-1 fibroblasts independently of their origin tend to reinstate their “normal/healthy” steady state (SS1). Fibroblasts from Cluster-3 tend to acquire a second steady state (SS2) which is distinctively different from the “normal” steady state. Blue dots indicate cluster number. (**D**) Cluster-2 fibroblasts can be divided into two sub-groups according to their velocity: one tending towards SS1 (C2G1, gold) and one towards SS2 (C2G2, magenta). (**E**) Histograms showing a ribosome score based on differentially expressed ribosomal protein genes in our dataset and a ribosome biogenesis score based on the expression of all GO genes associated with this process.

To further our understanding of the mechanisms driving disease onset, we calculated RNA velocity (22), which is based on the balance of unspliced (nascent) and spliced (mature) mRNA. This high-dimensional vector can act as an indicator of the future state of mature mRNA, driving cell state. The directionality of the plot arrows depicts the direction of cell state progression. RNA velocity analysis of all fibroblasts (**Figure 7C**) showed that the majority of the fibroblasts in Cluster-1 (composed mainly of non-IPF fibroblasts) trended towards a steady state (SS1). Even fibroblasts originating from IPF samples in this cluster appeared to be able to express genes driving them towards SS1. Similarly, cluster-3 fibroblasts are attracted to another steady state (SS2) which is likely disease related since almost all of these cells originated from IPF lungs. Although not as clear, Cluster-4 also showed a directional flow towards SS2 suggesting that these cells may transition to a profibrotic phenotype but have a cellular transcriptome that differentiates them from fibroblasts in Cluster-3 (**Figure 7C**). TIMP1, COL1A1, CTHRC1, TUBA1B, SH3BGRL3 and S100A16 were the top six differentially expressed genes between Cluster-3 and Cluster-4 (upregulated in Cluster-4).

Cluster-2 fibroblasts showed a more complex pattern (**Figure 7C**) with some cells going towards Cluster-1 (subgroup C2G1; **Figure 7D**, gold) and others towards Cluster-3 (subgroup C2G2; **Figure 7D**, purple) suggesting that cells in Cluster-2 were captured in a transition state that could either be resolved (SS1) or progress to a profibrotic state (SS2). We found 74 of the 103 genes coding for ribosomal proteins to be significantly downregulated in C2G2 vs C2G1 (**Table S5**). The remaining 29 ribosomal protein genes were unaffected. Based on these 74 genes, C2G1 and C2G2 cells were assigned a ribosome score which was significantly decreased in C2G2 (*p*-value << 0.001). In addition, we calculated a ribosome <u>biogenesis</u> score based on 267 other genes associated with this process (Gene Ontology Browser). Again, C2G2 showed a significant decrease for this score (*p*-value << 0.001). Both these observations (**Figure 7E**) are consistent with the hypothesis that there is a dysregulation of ribosomal function at the early stages of IPF.

To validate this finding, we used a second publicly available scRNA-seq dataset (6), that had similar tissue collection and processing procedures as ours. This dataset consisted of eight transplant donors and four IPF ex-plants, but it did not have distinct upper/lower lobe samples. We processed and reanalyzed the raw scRNA-seq data in the same way as our dataset. Due to the low number of IPF fibroblasts (see **Figure S4B)** we couldn’t confidently identify a subcluster similar to our Cluster-2 in the fibroblast cluster, but we observed three areas containing distinct cell populations originating from Controls, IPF and a mixture of the two, respectively (**Figure S4A**, green). Analysis of those cells showed 32 ribosomal genes to be differentially expressed between control and IPF with 29 of them downregulated. Notably, all 29 were also included in the previous list of 74 differentially expressed genes in our Cluster-2 (*p*-value<0.01).

### 3.3. Monooxygenase DBH Like 1 (MOXD1): a novel early IPF biomarker

To identify early IPF biomarkers specific to fibroblasts, we examined the Cluster-3 top upregulated genes that also showed low expression in the rest of the cell types (**Figure 5**). The five most significant of those included known IPF players (metalloprotease, collagen and two -LTBP1 LTBP2) and MOXD1 (**Table S3**). MOXD1 was not previously associated with IPF, but it is expressed significantly higher in Cluster 3 and it is significantly downregulated in controls (**Figure 8A-C**). Furthermore, it had been described in replicative senescent fibroblasts (27). Given that IPF fibroblasts can be resistant to apoptosis (28) and have a senescent phenotype (29), we selected MOXD1 for validation as a new possible fibroblast biomarker in early and advanced IPF.

**Figure 8.**
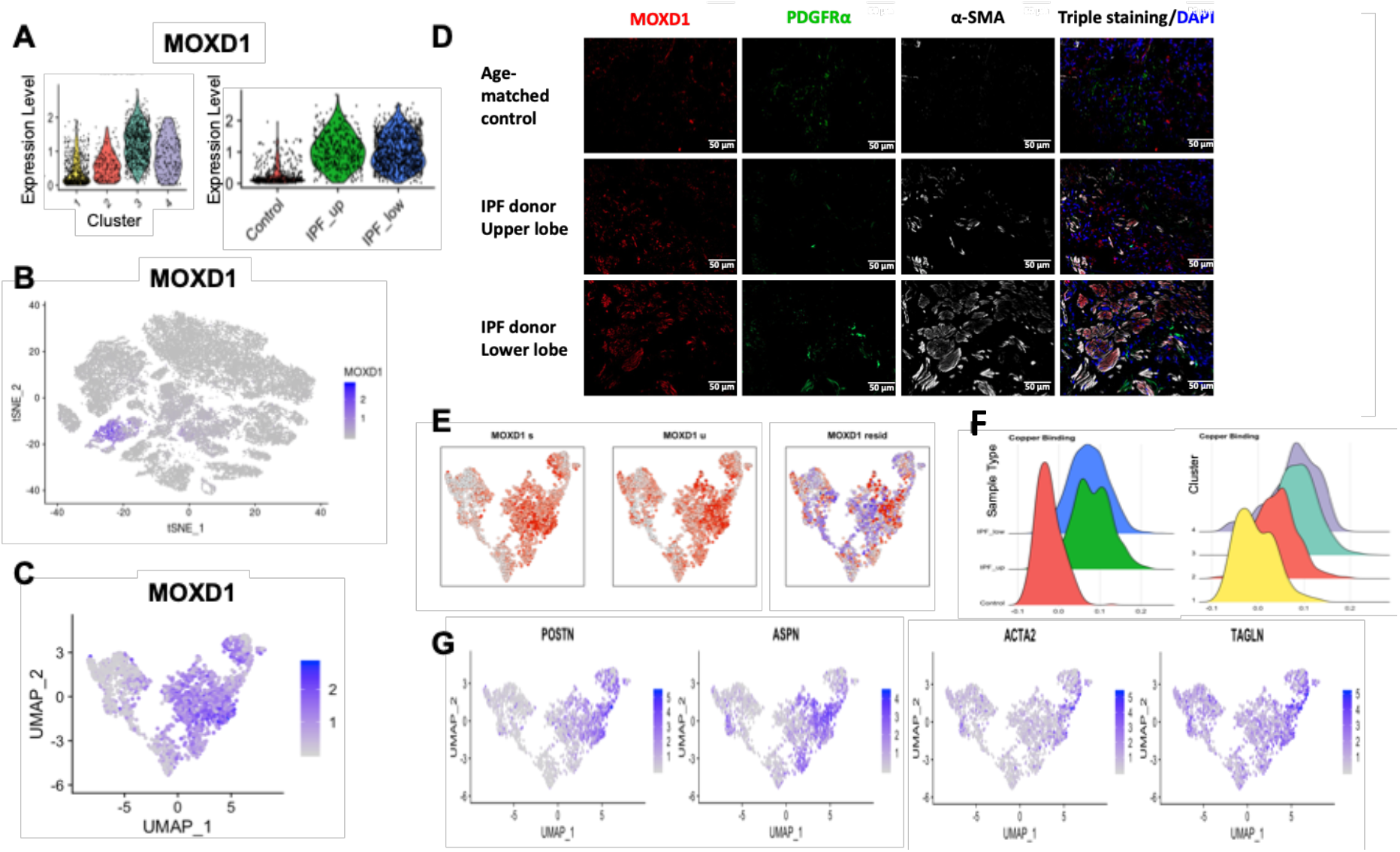
IPF fibroblasts express MOXD1. (**A**) MOXD1 expression peaks in Cluster-3 and is highly expressed in all IPF fibroblasts. (**B**) MOXD1 expression is specific for fibroblasts. (**C**) UMAP plot showing MOXD1 mRNA expression in all four fibroblasts clusters. (**D**) MOXD1 immunofluorescence staining in lung parenchyma frozen tissue, obtained from control and IPF lungs (x20 magnification, 50µm). *Red:* MOXD1; *green:* PDGFRα (fibroblast marker); *white*: α-SMA (myofibroblast marker). (**E**) Velocity analysis of MOXD1 mRNA. (**F**) Histograms showing the cell distribution based on a Copper-Binding-Score. (**G**) UMAP plots of myofibroblast expressing genes.

Immunofluorescence staining of human lung tissues (**Figure 8D** and **S5**) validated the mRNA findings. Cells from healthy donors’ tissues expressed minimal MOXD1 portein. IPF lung upper lobes, showing less advanced fibrotic disease, also showed increased MOXD1 compared to control lungs. IPF lower lobes, showing advanced fibrotic disease, expressed even higher levels of MOXD1. We then proceed to characterize the fibroblast cells expressing MOXD1 by co-staining the sections with PDGFRa (marker of fibroblasts) and alpha smooth muscle actin (α-SMA, a marker of myofibroblasts, which also marks pericytes and smooth muscle cells). As expected, α-SMA staining was increased in upper lobes and even more highly increased in lower lobes of IPF lungs compared to control lungs. Most MOXD1 staining in upper lobes was found in cells not staining with α-SMA, while these markers co-stained many of the same cells in the lower lobe suggesting that MOXD1 is a marker of cells that are maturing into myofibroblasts. This is consistent with increased levels of myofibroblasts in IPF. PDGFRa staining showed that fibroblasts in control lungs are mostly MOXD1/SMA negative, while fibroblasts in lower lobes of IPF lungs are mostly MOXD1/SMA positive Co-staining with these markers indicated that most of the MOXD1 staining cells in IPF are myofibroblasts To assess location of MOXD1 fibroblasts, we performed immunostaining of whole lung tissues. Lung sections from age matched donor control were negative for MOXD1, whereas sections from IPF lungs showed numerous positive cells (**Figure 8D**).

Velocity analysis of fibroblasts using their spliced (**Figure 8E**, left) and unspliced (**Figure 8E**, center) MOXD1 mRNA context indicated transcriptional kinetics towards the intersection between Cluster-3 and Cluster-4. A closer inspection of this region showed high expression of the myofibroblast associated genes POSTN, ASPN, ACTA2 and TAGLN (**Figure 8G**) (11, 30). It was clear from the UMAP plots that the cells carrying these myofibroblast markers did not segregate in one distinct cluster (**Figure 7C**) suggesting that myofibroblast specific genes are having similar signatures to the original cells that transitioned to a myofibroblast.

### 3.4. Copper-binding proteins are upregulated early in disease

MOXD1 is a copper-binding protein and copper itself has been implicated in pulmonary fibrosis (31). Blockhuys et al. (23) have identified 54 human Cu-binding proteins, divided in 9 groups according to their cellular localization. We found 39 of them to be significantly up- and 10 down-regulated in Cluster-3 (“fibrotic”) vs Cluster-1 (“control”) (FDR<0.05; **Table S6A**). Furthermore, 20 of them were already significantly upregulated in C2G2 compared to C2G1 subcluster (**Table S6B**). Among the top differentially expressed genes we found SPARC, LOXL1, and APP which have been implicated in lung fibrosis (32-34). In similar analysis to ribosome genes, we found 29 genes to be significantly upregulated in the validation cohort; and 24 of them were included in the 39 of Cluster-3. Next, we assigned a copper-binding score to all fibroblasts based on the expression of these proteins in our dataset. IPF samples, regardless of lung topology, showed higher scores compared to the control samples (**Figure 8F**, left). This difference was also reflected in the fibroblast clusters where Cluster-1 cells had the lowest Cu-binding score and Cluster-3 and 4 had the highest. Cluster-2, consistent with an intermediate cell state, showed an intermediate score (**Figure 8F**, right).

## 4. Discussion

Fibroblasts have been previously studied for their role in fibrosis in general and lung fibrosis in particular (35). Research has been focused on the lower subpleural lung regions where fibrotic scar tissue is primarily localized, assuming that fibroblasts from these regions contribute the most to disease development and progression. In the present work we are extending research to generally less disease-affected areas, the upper lobes of the lungs, to investigate early events in disease onset.

Our results show that the majority of IPF lung fibroblasts have a distinct phenotype. Interestingly, fibroblasts from the upper and lower lobes have similar molecular profiles; suggesting that gene expression changes in fibroblasts happen at an early disease stage. The observed early shift in gene expression in IPF fibroblasts, especially in extracellular matrix associated genes (collagens, MMP2, etc), agrees with computationally deconvoluted bulk RNA gene expression signatures (5). These models indicated that deciding regulatory molecular events can happen early or late in lung fibrosis, depending on the cell type. Our scRNA-seq results confirm this prediction.

One of the top differentially expressed genes in the IPF fibroblasts (Cluster-3) is MOXD1 which has not been previously studied in IPF. MOXD1 is a copper-binding enzyme, shows a tight association with the ER membrane and is not secreted (36). Currently its substrate is unknown. It is found to be upregulated in senescent human fibroblasts and human vascular endothelial cells (HUVEC) (27). Most importantly, IPF fibroblasts, under both *in vitro* and *in vivo* conditions and regardless of their lung localization, show high expression of MOXD1. Our velocity analysis results of MOXD1 nascent and mature mRNAs suggests that it start expressing earlier than known myofibroblast markers and could be potentially used for early diagnosis.

We find that MOXD1 upregulation in fibrotic cells is part of the more general upregulation of copper binding enzymes, including Lysyl Oxidase Like 1 (LOXL1) and Secreted Protein Acidic and Cystein Rich (SPARC), which have been studied for their pro-fibrotic role in multiple organs (34, 37). The dysregulation of copper-binding proteins along with the recent demonstration of copper induced lung fibrosis in mice (31) and copper-induced premature senescence of human fibroblasts (14) supports a more comprehensive role of copper and fibroblast senescence in the pathogenesis of IPF. It would be interesting to see whether IPF fibroblasts accumulate intracellular copper similar to the progressive accumulation of intracellular copper in aging, kidney fibrosis and other diseases (38-40).

Depending on their localization (IPF_LOW, IPF_UP), approximately 15% to 25% of fibroblasts from IPF samples retain a transcriptional profile that resembles that of normal control lung fibroblasts. However, they form a subgroup which indicates that their “normal” phenotype is influenced by the disease environment. Another 25% of IPF fibroblasts along with 15% of control fibroblasts seems to be in “perturbed” states that are characterized by increased expression of ECM genes. It is reasonable to assume that these states may represent normal wound healing processes occurring in both control and IPF lungs. In the case of IPF lungs, these processes would eventually run uncontrolled resulting in severe fibrosis (41).

One particular group of cells (Cluster-2) seem to capture this transition. It shows two distinct cell sub-populations with tendency towards opposing states. One subgroup shows a tendency to revert to a normal state while the other seems to be transitioning towards a profibrotic state. A closer examination of the profibrotic sub-population reveals a downregulation of almost all ribosomal proteins and dysfunctional ribosomal biosynthesis; and upregulation of the majority of copper-binding genes. We validate these findings in a similar cohort from a previous study (6). Posttranslational regulation has been shown to play an important role in heart fibrosis through the largely underexplored roles of RNA binding proteins and variation in ribosome occupancy, which affect protein expression levels independent of mRNA levels (42). Importantly, dysregulation of translational control on the level of polyribosome formation has been shown to play a significant role in the emergence of IPF myofibroblasts (43). Although we observe a decrease in ribosomal gene expression, we wouldn’t necessarily associate it with decreased genome-wide translational activity in these cells. Disproportionate expression of ribosomal proteins could affect ribosomal specificity (44, 45), favoring the translation of a profibrotic proteome in fibroblasts. Moreover, quantitative change in ribosome biogenesis could increase competition for ribosome binding and translation initiation, causing a variable effect on the translation of cellular mRNAs (46).

Impairment of ribosomal biogenesis not only could have an impact on the ribosomes’ cellular housekeeping role in protein synthesis but could also affect cell cycle and proliferation (15, 47). Ribosome biogenesis can be impaired at multiple steps by a variety of stress factors, resulting in ribosomal stress and cellular senescence (15-17). Our results support the hypothesis that ribosomal impairment may be a general mechanism activated during switching between two cell fates. In the future, experimental manipulation of ribosomal proteins will be necessary to decipher whether ribosome dysregulation is a driver of IPF or one of the many phenotypes associated with the dysregulated homeostasis caused by the disease.

In summary, we examine the events that take place in fibroblasts when cells transition from healthy state to disease. We demonstrate that, at the single cell level, changes in the transcriptome of fibroblasts appear early in the relatively unaffected upper lobes of the IPF lungs. We also identify two focal points (steady states) towards where the normal and the fibrotic cells coalesce. More importantly, we discover a point of transition between these two states, which is characterized by a significant decrease in ribosomal protein genes and a significant increase in copper-binding proteins. We examine the second most upregulated copper-binding protein, MOXD1, and find it to be fibroblast- and IPF-specific. Furthermore, we find that MOXD1 is also expressed at early stages of IPF suggesting that it can be used as an early biomarker of fibroblast pulmonary fibrosis.

Increasing evidence supports the idea that IPF fibroblasts have acquired resistance to apoptosis (28) and have a senescent phenotype (29). Furthermore, senescence can mediate the fibrotic phenotype (48), perhaps by secreting profibrotic factors (49). Thus, early targeting of senescent fibroblasts with senolytic cocktails may help manage or even reverse the disease (48, 49). To that extend, in future studies the transitional fibroblasts we discovered can farther our mechanistic understanding of the transition process and can be utilized for the development of potential diagnostic and therapeutic strategies.

At present, a major limitation of this study is the lack of available biomarkers for these transitioning fibroblasts. Future studies can reveal how senescence and alteration of ribosomal protein and copper-binding pathways may affect the switch of fibroblasts from the normal steady state to a profibrotic state. Furthermore, in-depth studies of these fibroblasts may provide a useful diagnostic tool for early detection of IPF and possibly a therapeutic window before extensive tissue damage occurs.

## DECLARATIONS

### Ethics statement

All data obtained from previously published samples from our work or work of others.

### Source of materials

All data presented in this manuscript are available from GEO (GSE128033, GSE122960).

### Competing interests

RL served as a consultant for Bristol Myers Squibb, Formation, Sanofi, Biocon, Boehringer-Mannheim, Merck and Genentech/Roche, and holds or recently had research grants from Corbus, Formation, Elpidera, Regeneron, Pfizer and Kiniksa.

### Funding

This work was funded by the National Institutes of Health grant U01HL145550 to PVB, RL, ALM, MR; and U01HL137159 and R01LM012087 to PVB.

### Authors’ contributions

MJ: study design, performed data analysis, interpreted the results, wrote the paper. LR, TC and AB: collected data (tissue stainings), interpreted the results. MGK: interpreted the results, wrote the paper. TT: performed data analysis. JS: collected tissues, performed experiments. ALM: interpreted the results, helped with paper writing. RL: interpreted the results, helped with paper writing. MR: interpreted the results, helped with paper writing. PVB: study design, interpreted the results, wrote the paper.

## Notes

https://www.ncbi.nlm.nih.gov/geo/query/acc.cgi?acc=GSE128033

https://www.ncbi.nlm.nih.gov/geo/query/acc.cgi?acc=GSE122960

