## Supplementary material for "Early events marking lung fibroblast transition to profibrotic state in idiopathic pulmonary fibrosis": Suplpemental Figures

### SUPPLEMENTAl FIGURES

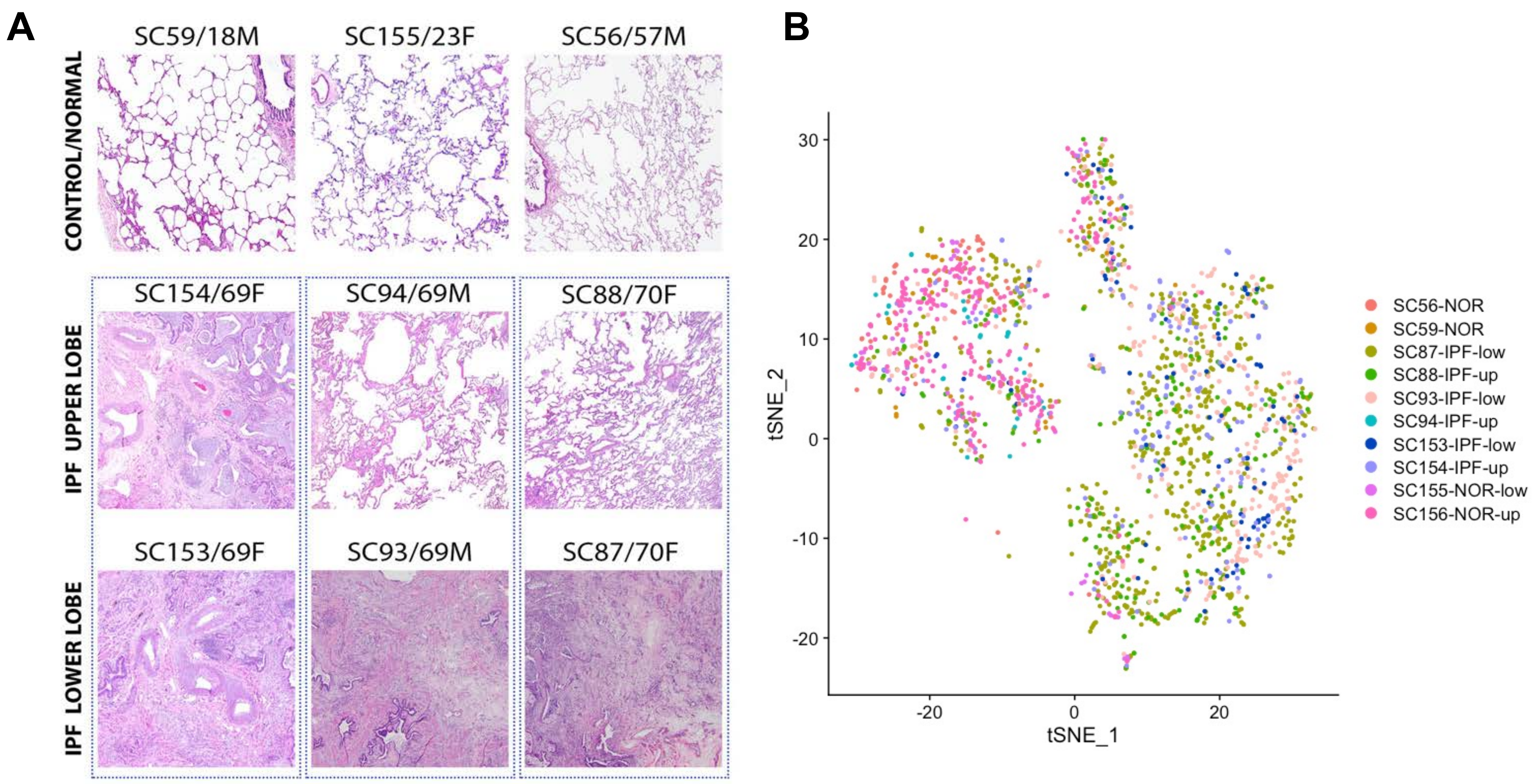
A
SC59/18M
SC155/23F
SC56/57M
CONTROL/NORMAL
SC154/69F
SC94/69M
SC88/70F
IPF UPPER LOBE
SC153/69F
SC93/69M
SC87/70F
IPF LOWER LOBE
B
30
20
10
0
-10
-20
tSNE_2
-20
0
20
tSNE_1
SC56-NOR
SC59-NOR
SC87-IPF-low
SC88-IPF-up
SC93-IPF-low
SC94-IPF-up
SC153-IPF-low
SC154-IPF-up
SC155-NOR-low
SC156-NOR-up

**FIGURE S1**

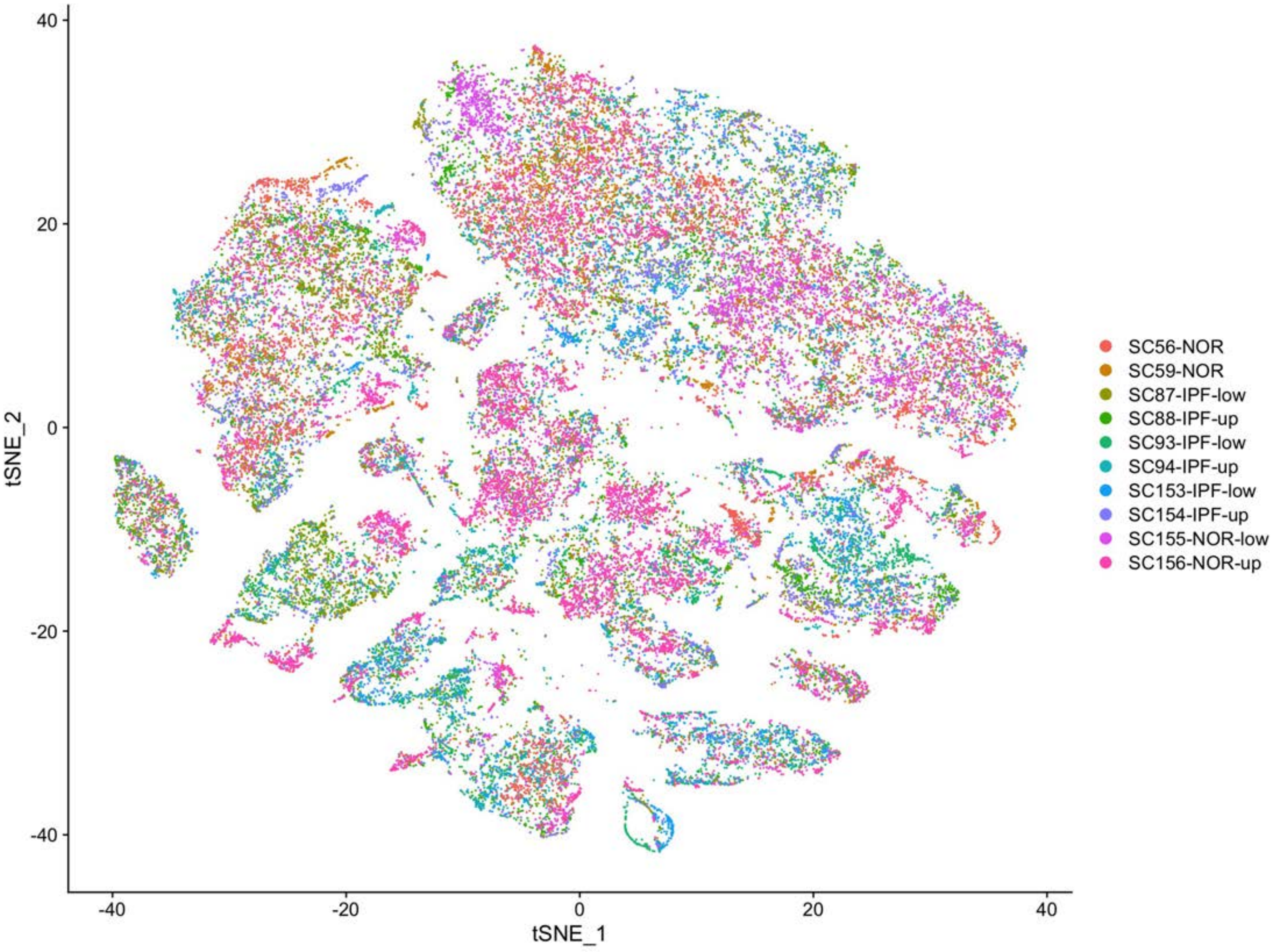

40
20
0
-20
-40
tSNE_2
-40
-20
0
20
40
tSNE_1
SC56-NOR
SC59-NOR
SC87-IPF-low
SC88-IPF-up
SC93-IPF-low
SC94-IPF-up
SC153-IPF-low
SC154-IPF-up
SC155-NOR-low
SC156-NOR-up


**FIGURE S2**

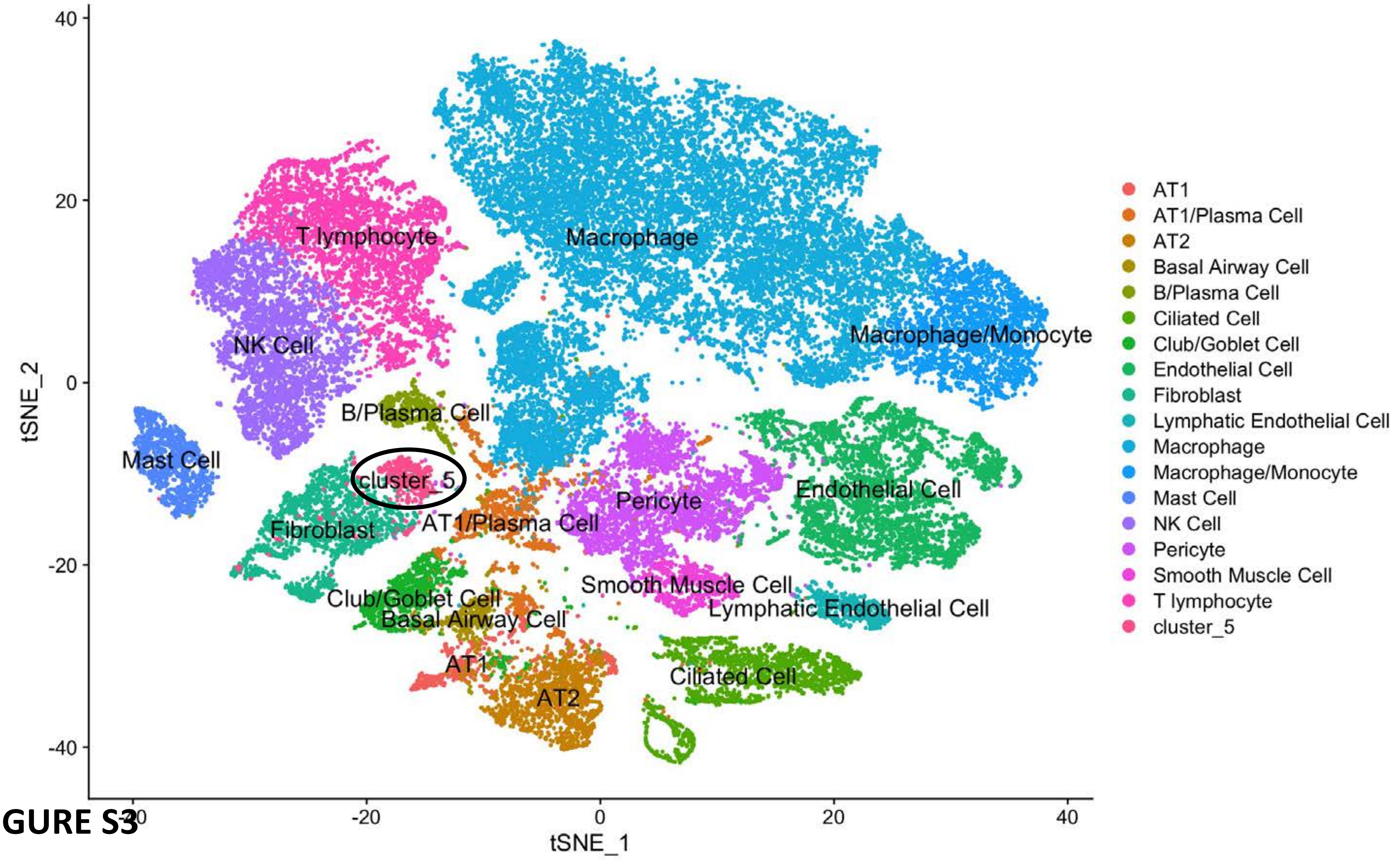

40
20
0
-20
-40
tSNE_2
-20
0
20
40
tSNE_1
T lymphocyte
Macrophage
NK Cell
Macrophage/Monocyte
B/Plasma Cell
Mast Cell
cluster_5
Pericyte
Endothelial Cell
Fibroblast
AT1/Plasma Cell
Smooth Muscle Cell
Club/Goblet Cell
Lymphatic Endothelial Cell
Basal Airway Cell
AT1
Ciliated Cell
AT2
AT1
AT1/Plasma Cell
AT2
Basal Airway Cell
B/Plasma Cell
Ciliated Cell
Club/Goblet Cell
Endothelial Cell
Fibroblast
Lymphatic Endothelial Cell
Macrophage
Macrophage/Monocyte
Mast Cell
NK Cell
Pericyte
Smooth Muscle Cell
T lymphocyte
cluster_5


**FIGURE S3**

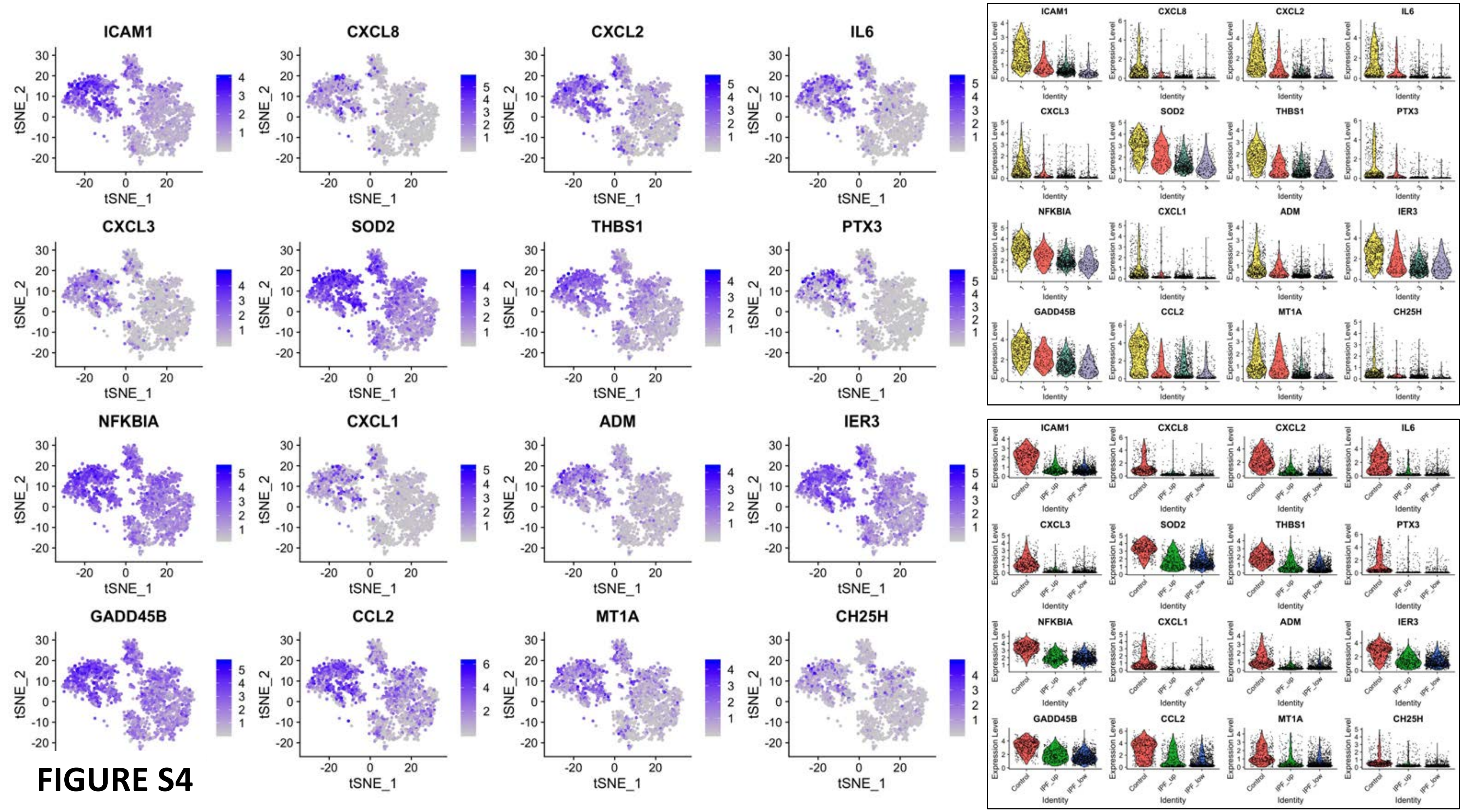
ICAM1
CXCL8
CXCL2
IL6
CXCL3
SOD2
THBS1
PTX3
NFKBIA
CXCL1
ADM
IER3
GADD45B
CCL2
MT1A
CH25H
tSNE_1
tSNE_2
Expression Level
Identity
Control
IPF_up
IPF_low

FIGURE S4

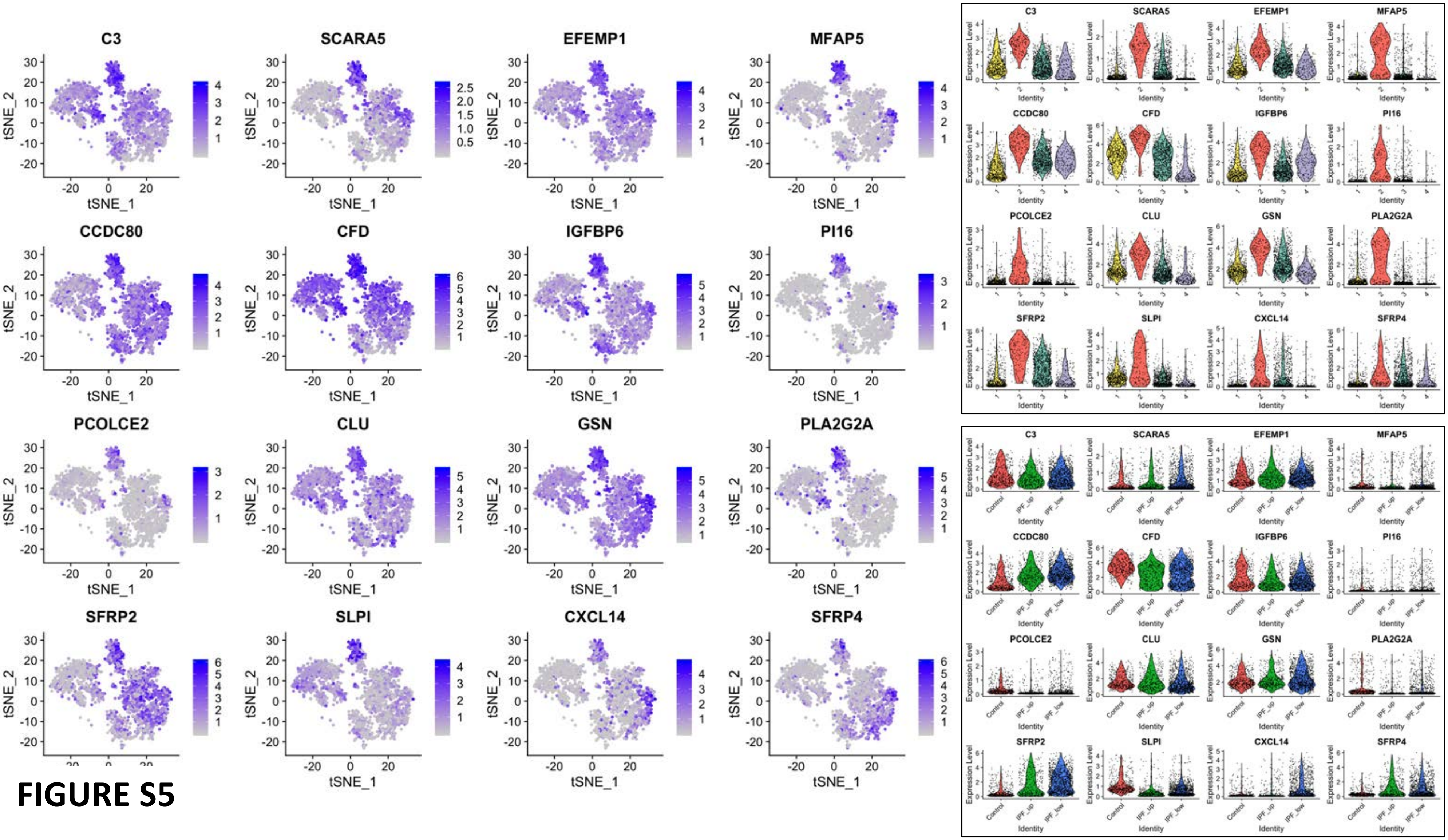
C3
SCARA5
EFEMP1
MFAP5
CCDC80
CFD
IGFBP6
PI16
PCOLCE2
CLU
GSN
PLA2G2A
SFRP2
SLPI
CXCL14
SFRP4
tSNE_1
tSNE_2
Expression Level
Identity
Control
IPF_up
IPF_low

FIGURE S5

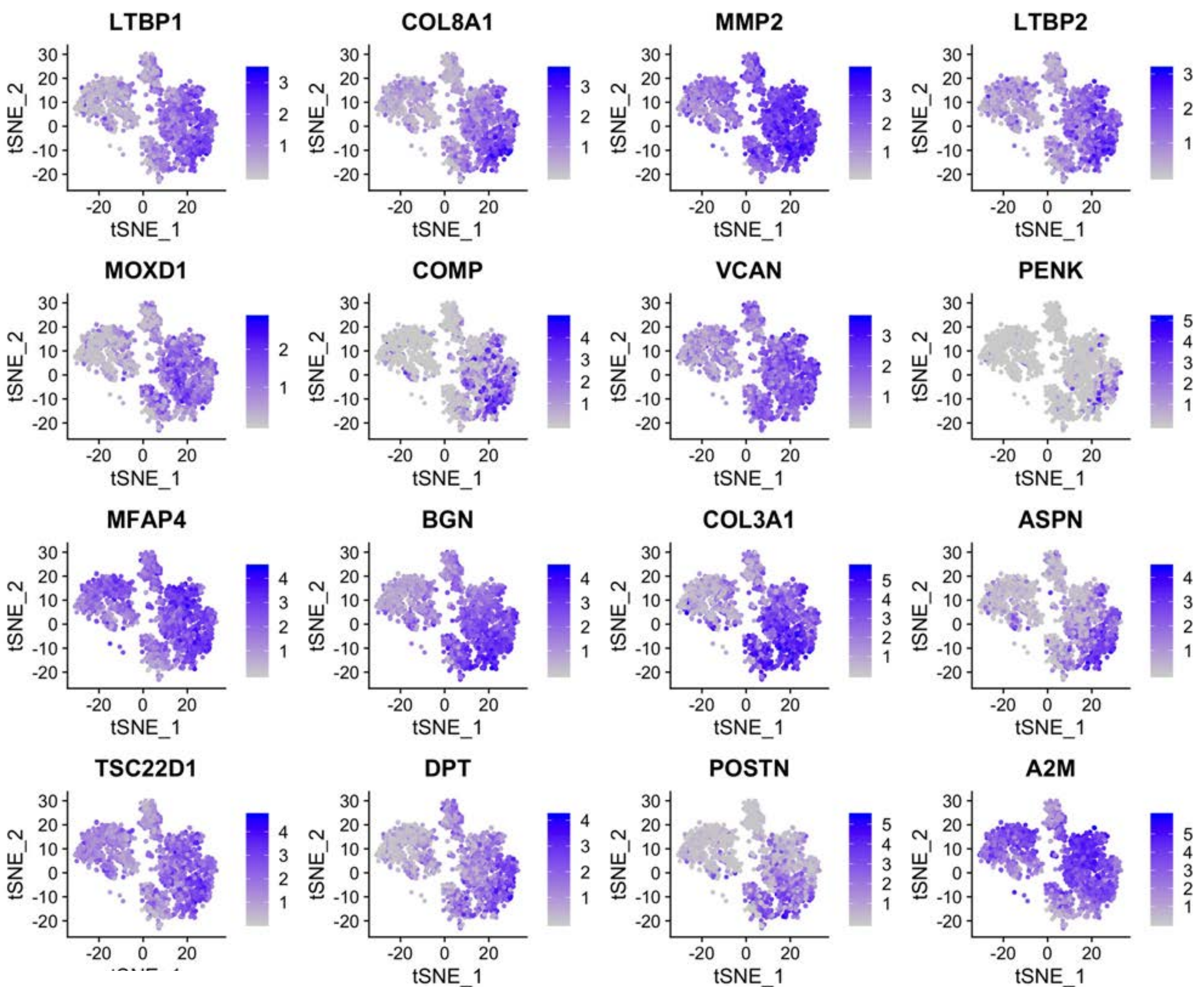
LTBP1
COL8A1
MMP2
LTBP2
MOXD1
COMP
VCAN
PENK
MFAP4
BGN
COL3A1
ASPN
TSC22D1
DPT
POSTN
A2M
tSNE_1
tSNE_2

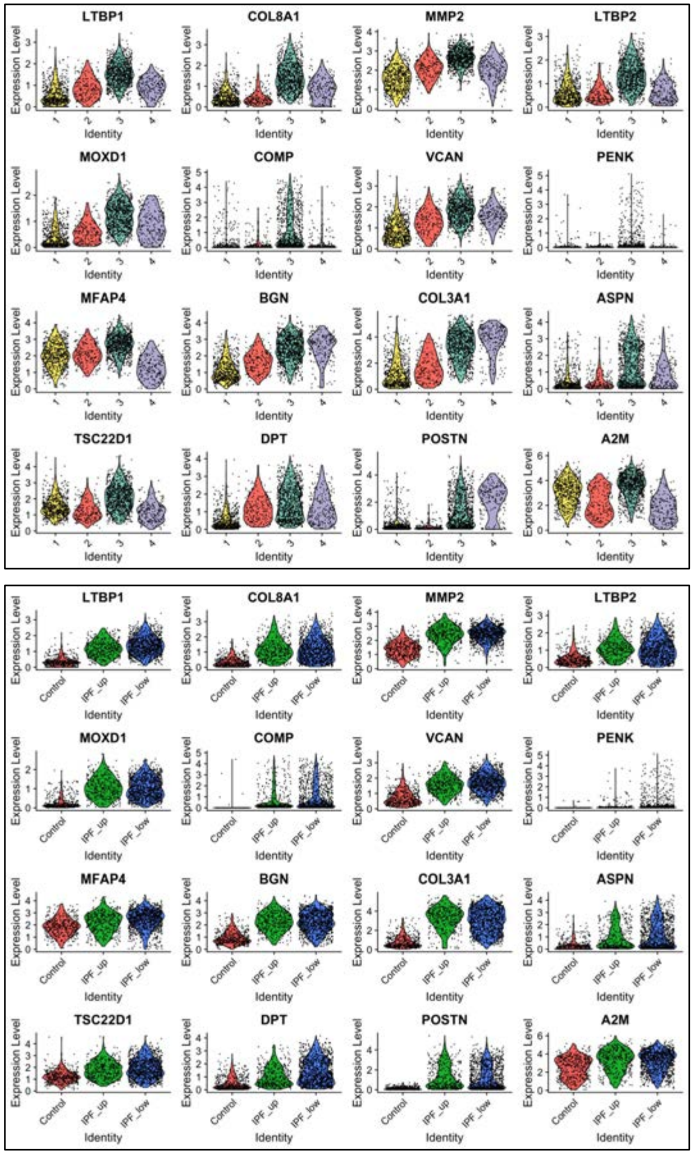
LTBP1
COL8A1
MMP2
LTBP2
MOXD1
COMP
VCAN
PENK
MFAP4
BGN
COL3A1
ASPN
TSC22D1
DPT
POSTN
A2M
Expression Level
Identity
Control
IPF_up
IPF_low

**FIGURE S6**

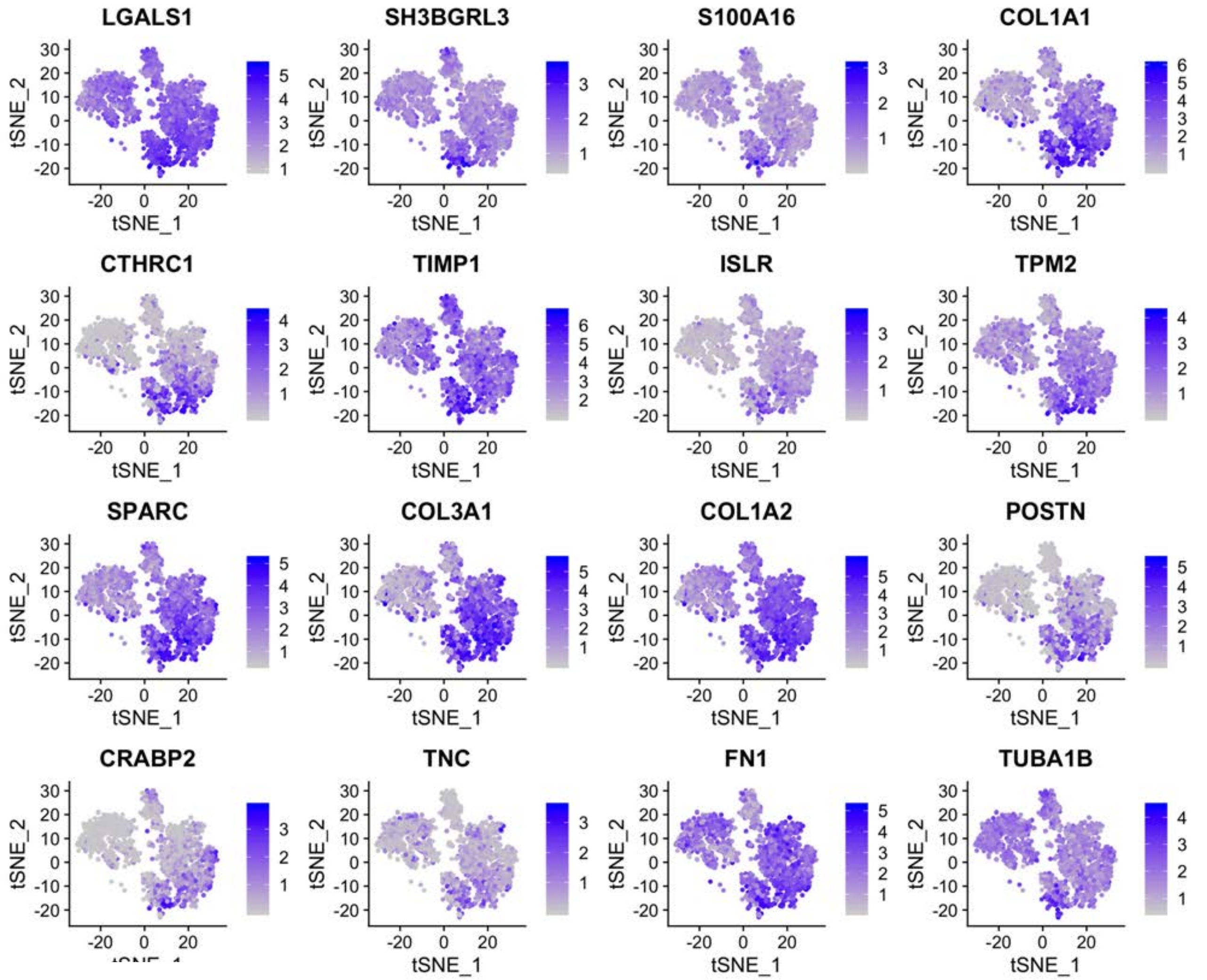
LGALS1
SH3BGRL3
S100A16
COL1A1
CTHRC1
TIMP1
ISLR
TPM2
SPARC
COL3A1
COL1A2
POSTN
CRABP2
TNC
FN1
TUBA1B
tSNE_1
tSNE_2

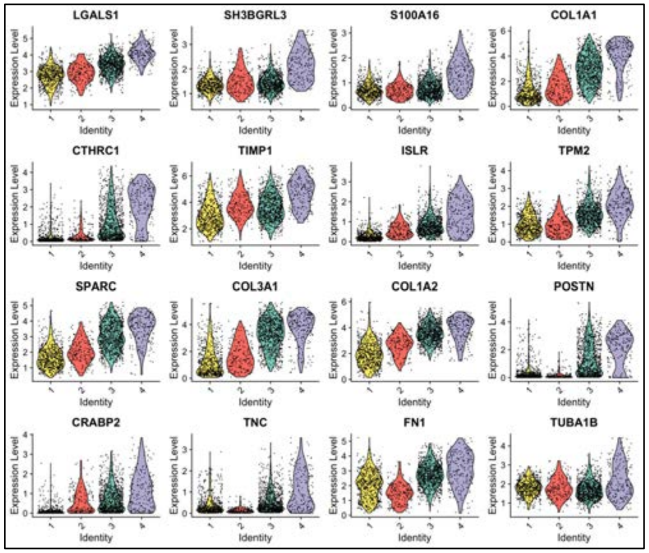

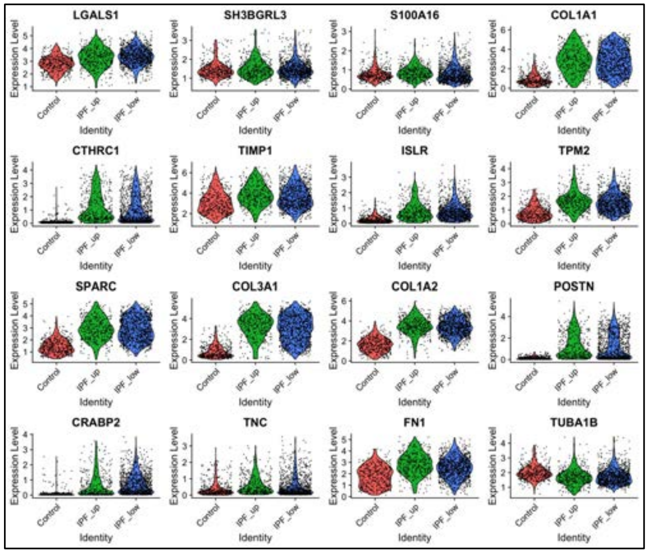

**FIGURE S7**

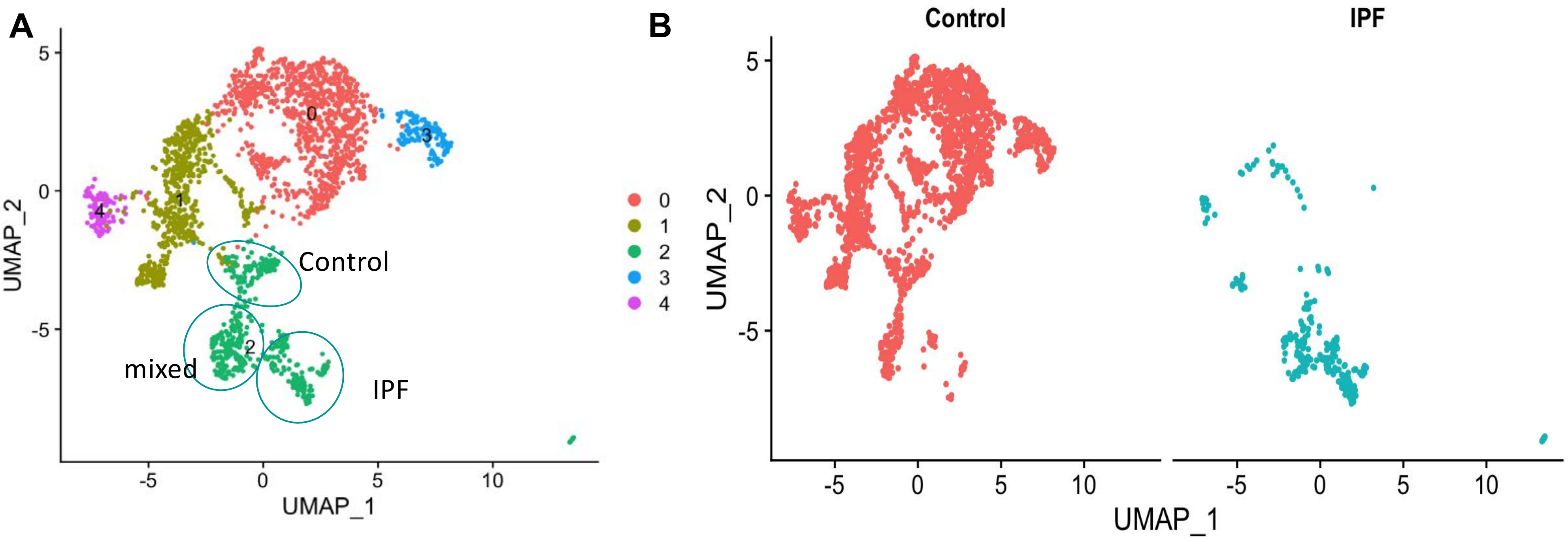


**FIGURE S8**

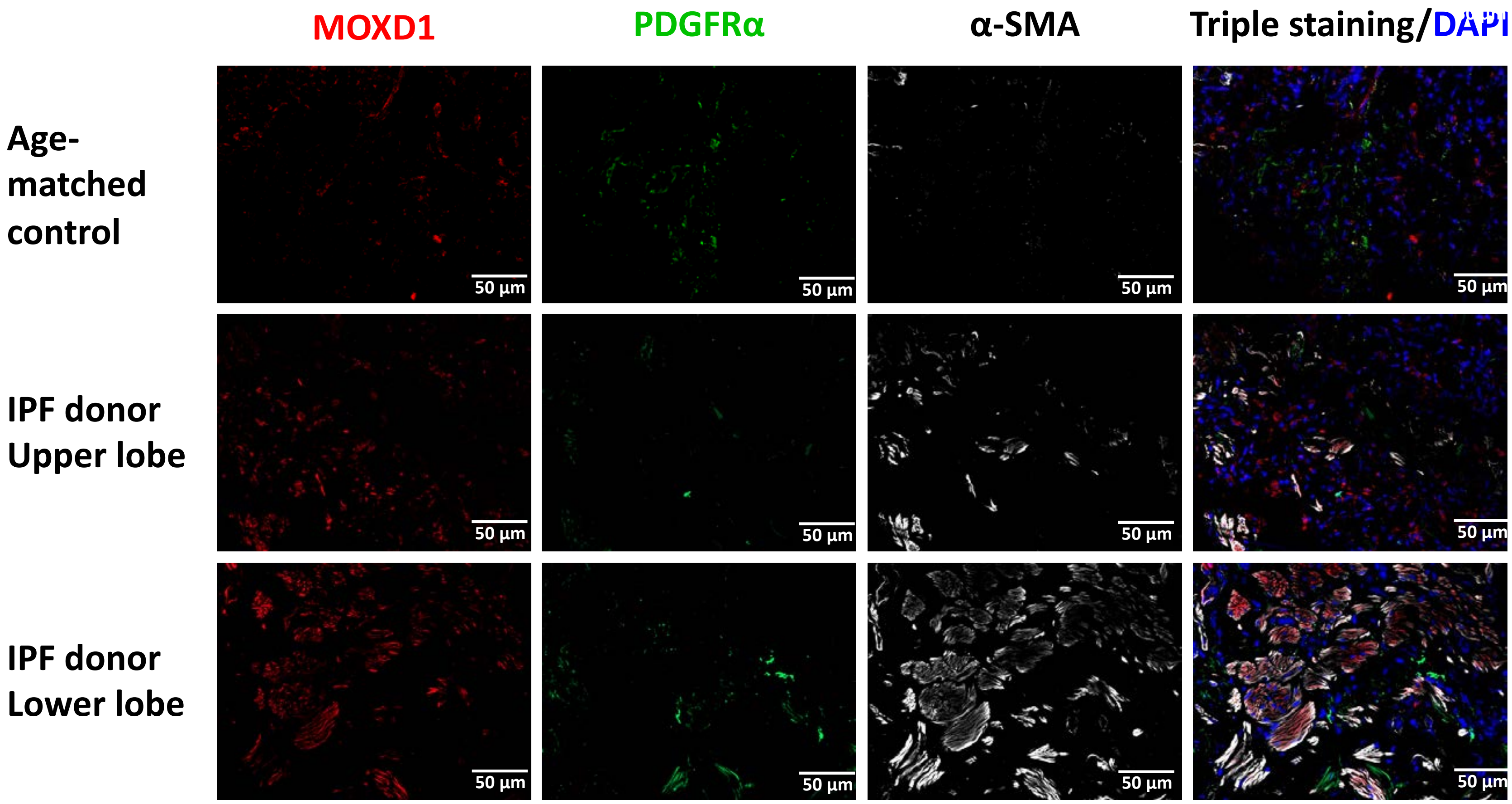
MOXD1
PDGFRα
α-SMA
Triple staining/DAPI
Age-matched control
IPF donor Upper lobe
IPF donor Lower lobe
50 µm

### FIGURE S9

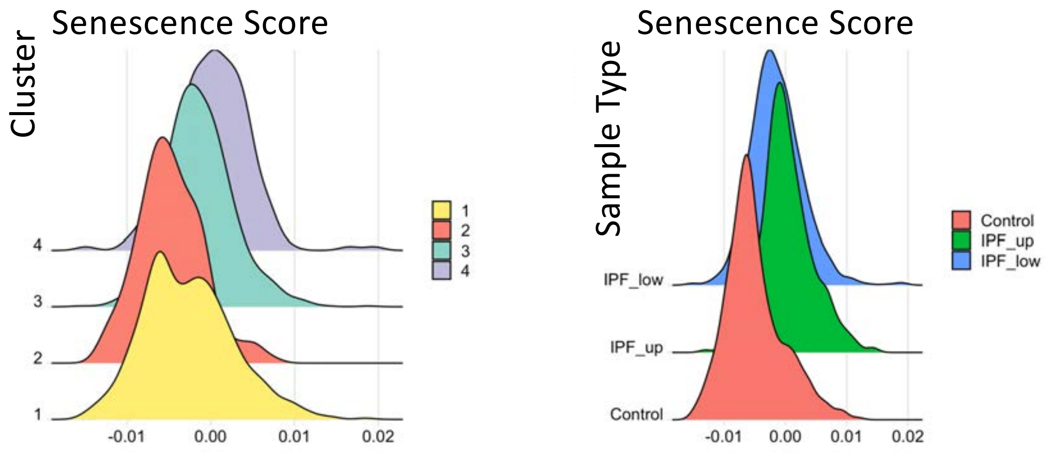
Senescence Score
Cluster
4
3
2
1
-0.01
0.00
0.01
0.02
1
2
3
4
Senescence Score
Sample Type
IPF_low
IPF_up
Control
-0.01
0.00
0.01
0.02
Control
IPF_up
IPF_low

**FIGURE S10**
